# *Iba1+* Microglia Exhibit Morphological Differences between Inferior Colliculus Sub-Regions and Their Abutments onto *GAD67+* Somata Reveal Two Novel Sub-types of GABAergic Neuron

**DOI:** 10.1101/606509

**Authors:** Samuel David Webb, Llwyd David Orton

**Affiliations:** Department of Life Sciences, Manchester Metropolitan University, Manchester, M1 5GD; Institute of Neuroscience, Newcastle University, Newcastle upon Tyne, NE2 4HH, UK

**Keywords:** Auditory, Colliculi, Glia, Anatomy, Confocal microscopy, Immunohistochemistry

## Abstract

Microglia have classically been viewed as the endogenous phagocytes of the brain, however, emerging evidence suggests roles for microglia in the healthy, mature nervous system. We know little of the contribution microglia make to ongoing processing in sensory systems. To explore *Iba1+* microglial diversity, we employed the inferior colliculi (IC) as model nuclei, as they are characterized by sub-regions specialized for differing aspects of auditory processing. We conducted fluorescent multi-channel immunohistochemistry and confocal microscopy in guinea pigs of both sexes and discovered that the density and morphology of *Iba1+* labelling varied between parenchymal sub-regions of IC, while *GFAP+* labelling of astrocytes was confined to the *glia limitans externa* and *peri*-vascular regions. The density of *Iba1+* microglia somata was similar across sub-regions, however a greater amount of labelling was found in dorsal cortex than ventral central nucleus or lateral cortex. To further understand these differences between sub-regions in IC, Sholl and skeleton analyses of individual microglia revealed a greater number of branching ramifications in dorsal cortex. We also quantified abutments of *Iba1+* microglial processes onto *GAD67+* (putative GABAergic) somata. Cluster analyses revealed two novel sub-types of *GAD67+* neuron, which can be distinguished solely based on the quantity of axo-somatic *Iba1+* abutments they receive. These data demonstrate *Iba1+* microglia exhibit different morphologies and interactions with *GAD67+* neurons in distinct sub-regions of the mature, healthy IC. Taken together, these findings suggest significant heterogeneity amongst microglia in the auditory system, possibly related to the ongoing functional demands of their niche.

## Introduction

Inhibition is an essential element of neural processing and a defining component of sensory systems. GABAergic inhibition is prevalent in the auditory system, particularly in the principal auditory midbrain nuclei, the inferior colliculi (IC). Around a quarter of neurons in IC are GABAergic (Merchán et al., 2005), which may be why the IC is the most metabolically active nucleus in the mammalian brain (Sokoloff et al., 1977). Understanding how inhibitory cell types vary in different brain regions, to specialize for distinct functions, is a key area of neuroscientific study (Freund and Buzsaki, 1996; Tremblay et al., 2016). Most investigations into sub-types of inhibitory neurons naturally focus on the cells *per se*, including their morphology, electrophysiological firing characteristics, expression of cytoplasmic calcium binding proteins and RNA transcriptome. Another approach is to characterize and classify GABAergic neurons based on differences in the afferent axo-somatic inputs they receive (Ito et al., 2009; Beebe et al., 2016).

The IC has a tonotopic topography that can be divided into distinct sub-regions. The central nucleus (CNIC) is dominated by neurons sharply tuned to simple auditory stimuli. The dorsal cortex (DCIC) has much broader frequency tuning and receives extensive corticofugal input and is specialized for synaptic plasticity (Herbert et al., 1991; Winer et al., 1998; Bajo and Moore, 2005; Bajo et al., 2010). The other major sub-region is the lateral cortex (LCIC) which exhibits polysensory tuning (Aitkin et al., 1978). Despite the essential role of the IC in hearing, little is known of how glial cells contribute to processing therein.

Microglia are an integral cell type present throughout the brain. The morphology of microglia varies throughout the brain, suggesting adaptation to their surrounding milieu (Lawson et al., 1990). Furthermore, microglia interact with neurons during ‘normal’ processing and can sense and respond to local chemical signaling (Pocock and Kettenmann, 2007; Wake et al., 2009; Schafer et al., 2012), but these processes remain poorly understood.

Here, we take advantage of the functional organization of IC sub-nuclei to investigate the anatomical inter-relationships of *Iba1+* microglia and *GAD67+* (putative GABAergic) neurons. We employed multi-channel fluorescence immunohistochemistry and confocal microscopy, and tested the hypothesis that microglia would form sub-region specific adaptations to their niche.

## Materials and methods

### Regulation and Ethics

All animals were housed and procedures performed in accordance with the terms and conditions of a license (PPL 60/3934) issued by the UK Home Office under the Animals (Scientific Procedures) Act 1986. Ethical approval was provided by the Local Ethical Review committee at Newcastle University and the School of Healthcare Science Ethics Committee at Manchester Metropolitan University.

### Animals

Results are described from four adult (one at four months old, three at six months old) outbred, tricolor guinea pigs (*Cavia porcellus*) of both sexes (three male, one female). Animal weights on the morning of each respective perfusion ranged from 675g to 867g. We aimed to minimize the number of animals used and their suffering at all times.

### Anesthesia and Tissue Processing

Animals were deeply anesthetized with sodium pentobarbital (i.p. injection; Euthanal, Merial; 200mg/ml, 2 ml volume). After five minutes, the pedal withdrawal and blink reflexes were assessed to confirm a deep plane of anesthesia. This was followed by transcardial gravity perfusion with 500mls of 0.1M heparinized PBS followed by 500mls of freshly made 4% paraformaldehyde in 0.1M PBS. Both solutions were pH 7.2 directly prior to use.

Brains were removed with rongeurs (Micro Friedman, 0.8mm jaws, WPI) and post-fixed in 30% sucrose in 4% paraformaldehyde, for at least 3 days at 4°C. Once brains sank, they were cut in the coronal plane with a razor blade through the parietal and temporal lobes, at around the rostro-caudal location of the medial geniculate. The tissue block was then placed in an embedding mold (Peel-a-way; Shandon), covered in embedding medium (OCT; Agar Scientific) and frozen at −80°C. 60µm sections were taken on a cryostat (HM560, Microm) and collected in 12 well plates in cryoprotectant (30% sucrose, 30% ethylene glycol, 1% polyvinyl pyrrolidone-40 in 0.1M PBS) and stored at −20°C until use (Watson et al., 1986; Olthof et al., 2019).

### Antibody characterization

The following primary antibodies were used:

*Mouse anti-GAD67* (1:500; monoclonal; clone 1G10.2; MAB5406; lot# 2636700; Millipore; RRID: AB_2278725) – according to the manufacturer, the immunizing antigen is a recombinant fusion protein containing unique N-terminus regions from amino acids 1-101 of *GAD67*. Immunoblotting detects a 67kDa protein in rat cerebellum and mouse microsomes; immunohistochemistry demonstrated labelling similar in distribution to *in situ* mRNA hybridization (Fong et al., 2005; Kotti et al., 2006; Ramirez et al., 2008). Use of this antibody has been published in guinea pig IC (Nakamoto et al., 2013; Foster et al., 2014; Beebe et al., 2016), as well as rat IC (Ito et al., 2009).

*Mouse anti-GFAP* (1:500; monoclonal; clone G-A-5; G3893; lot# 045M4889V; Sigma; RRID: AB_477010) – according to the manufacturer, this antibody is raised against an epitope from the C-terminus of *GFAP* in purified pig spinal cord (Latov et al., 1979; Debus et al., 1983). The antibody has been shown to recognize a single band of approximately 50kDa and reacts with homologous, conserved residues across mammals (Lorenz et al., 2005). The use of this antibody has been demonstrated in many species, including mouse (Komitova et al., 2005), rat (Lennerz et al., 2008; Sanchez et al., 2009), tree shrew (Knabe et al., 2008), guinea pig (Kelleher et al., 2011; Kelleher et al., 2013) and human (Toro et al., 2006). Labelling observed in this study was consistent with these studies and the known morphology of astrocytes.

*Rabbit anti-calbindin D-28k* (1:1,000; polyclonal; AB1778; lot# 2895780; Millipore; RRID: AB_2068336) – according to the manufacturer, this antibody recognizes a single band at 28kDa in human, mouse, and rat brain tissues. It does not bind to calretinin and pre-adsorbtion of diluted antiserum with calbindin removed all labelling in human brain (Huynh et al., 2000). Previous labelling of mouse olfactory bulb (Kotani et al., 2010), rat piriform cortex (Gavrilovici et al., 2010) and guinea pig enteric nervous system (Liu et al., 2005) all showed highly selective cytoplasmic labelling of neurons. We observed labelling consistent with previous reports.

*Rabbit anti-calretinin* (1:1,000; polyclonal; AB5054; lot# 2903043; Millipore; RRID: AB_2068506) – according to the manufacturer, this antibody recognizes the 29kDa protein in mouse and rat tissues (Su et al., 2010; Yanpallewar et al., 2010). This highly conserved epitope has also been labelled in hamster (Lee et al., 2004), zebrafish (Goodings et al., 2017) and turtle (Parks et al., 2017). We observed labelling consistent with these previous reports.

*Rabbit anti-Iba1* (1:1,000; polyclonal; 019-19741; lot# WDE1198; Wako; RRID: AB_839504) – according to the manufacturer, this affinity purified antibody was raised against a synthetic peptide corresponding to the C-terminus fragment of rat *Iba1*. Labelling via western blot was positive for a 17kDa band (Imai et al., 1996). We observed selective labelling of ramified microglia, matching similar reports in mouse (Bulloch et al., 2008), rat (Helfer et al., 2009; Fuentes-Santamaría et al., 2012), Japanese quail (Mouriec and Balthazart, 2013), macaque (Stanton et al., 2015) and chimpanzee (Rosen et al., 2008). We observed labelling consistent with these previous reports.

Blood vessels were labelled with rhodamine conjugated Griffonia (Bandeiraea) Simplicifolia Lectin 1 (1:100; RL-1102; Vector; lot# S0926RRID: AB_2336492), which binds to glycoproteins lining the inner lumen.

### Fluorescence immunohistochemistry

Sections through the superior colliculus and the rostral-most third along the rostro-caudal axis through the IC were first used to optimize labelling. Data are presented from sections in the middle third of the IC along the rostro-caudal axis, which contained the CNIC, DCIC and LCIC. The location of each section through the rostro-caudal axis was referenced to an atlas of the guinea pig brainstem (Voitenko and Marlinsky, 1993).

All steps in the labelling protocol involved continuous gentle agitation of sections. Free-floating sections were brought to room temperature and washed 3×5mins in PBS. Sections were blocked and permeabilized in 5% normal goat serum (Vector) and 0.05% Triton X-100 (Sigma) in PBS for one hour. Following blocking, a cocktail of primary antibodies was added to the blocking solution and applied to sections overnight at room temperature. The next day, sections were washed 3×5mins in PBS and incubated for two hours in appropriate secondary antibodies (Invitrogen; 1:250 in blocking solution). For double labelling of *Iba1* and *GAD67*, goat anti-rabbit AlexaFluor 488 and goat anti-mouse AlexaFluor 568 were used. For double labelling of calbindin and *GFAP*, goat anti-rabbit AlexaFluor 488 and goat anti-mouse AlexaFluor 647 were used. For triple labelling of *Iba1*, *GSL1* (pre-conjugated rhodamine fluorophore) and *GFAP*, goat anti-rabbit AlexaFluor 488 and goat anti-mouse AlexaFluor 647 were used. Sections were then mounted on slides and coverslipped using Vectashield (Vector Labs, H-1000) and kept at 4°C until imaged. All experiments had control slides where the primary, secondary or both the primary and secondary antibodies were excluded. This allowed detection of autofluoresence and any aspecific signal and ensured only labelling from primary and secondary binding to epitope targets was imaged.

### Image acquisition

Sequentially acquired micrographs were taken with a confocal microscope (Leica SP5) using a wide field stage and zoom function. Images were acquired via a 40x objective (NA=1.25) for images of the entire cross-section of the IC, and a 63x objective (NA=1.4) for region of interest (ROI) panoramas. Whole IC images were taken using 5µm equidistant slices in the Z-plane to produce maximum intensity tiled projections (pixel size; x & y=0.7583µm, z=50-60µm). For *GAD67* and *Iba1* ROI panoramas, 5-row × 6-column (432×552µm) tiled images were taken using 1µm z-slices and rendered as maximum intensity projections (pixel size; x & y=0.2406µm, z=40µm).

### Image analyses

For *Iba1+* cell density estimates, tiled panorama images of the IC were subject to manual cell counts. The peripheral borders of the IC were delineated and a contour drawn, and each image cropped to its respective contour. To make fair comparisons between cases, 450µm^2^ grids were placed across each IC panorama image and centered on the middle pixels of each micrograph in ImageJ. Only those grids which were filled entirely by stained parenchymal tissue were subject to counts. Comparisons were then made between cases, such that only grids that were present in images from all four animals were included in calculation of group means and standard deviations per grid.

Maximum intensity projection ROI panoramas were analyzed for i) cell counts, ii) percentage field of view covered analyses, iii) individual *Iba1+* cell Sholl analyses, iv) skeleton analyses and v) *Iba1* abutting *GAD67* analyses using Fiji ImageJ (Abràmoff et al., 2004). For analyses i-iv, panorama micrographs were first processed by filtering monochrome images using a median pixel (1.5) filter and then thresholded to binary by implementing the IsoData algorithm. Cell counts, and percentage field of view covered analyses were then performed using the Analyze Particles plugin. For Sholl analyses, individual *Iba1+* microglia were cropped and a series of equidistant radiating 1µm concentric circles were plotted from the center of the cell body to the furthest radiating extent of ramification. Each intersection with a concentric ring was measured. The Skeletonize algorithm was used to display a one-pixel thick framework of each microglial cell. The Analyze Skeleton plugin calculated number of branches and branch lengths for each cell.

Five *GAD67+* cells were selected randomly from each sub region panorama, with a selection criterion that the entirety of the cell must be contained within the x, y and z dimensions of the field of view. A z-stack was collected beyond the limits of each cell using 1µm slices. The absence or presence of the soma in each slice was determined and these data were used to calculate soma diameter to the nearest µm. Soma perimeters were manually contoured in each slice of each *GAD67+* cell. *Iba1+* labelling that came into contact with the contour with no pixels between was counted as an *Iba1+* process abutting a *GAD67+* cell soma. These data were calculated for each *Iba1+* cell as well as each *GAD67+* cell. The length of *Iba1+* labelling abutting the perimeter contour around each *GAD67+* cell soma was measured.

### Statistical analysis

Data were collected in Excel spreadsheets. Statistical hypothesis testing was performed in Prism 7 (GraphPad). Factorial analyses were conducted using the non-parametric Kruskall-Wallis ANOVA with sub-region as the factor in all cases. Where appropriate, *post-hoc* tests with Dunn’s method were conducted. For *post-hoc* analyses the α was Šidák corrected for multiple comparisons. Spearman’s rank correlations were used to investigate potential associations between dependent variables.

Principal component and two-step cluster analyses were conducted in SPSS v25 (IBM). The two-step cluster analysis employed Euclidean distance measures with Schwarz’s Bayesian clustering criterion and classified data into one of the two identified clusters. All 160 cell ROIs were successfully classified by this analysis. A Chi-squared test was used to analyze the ratio of cells in IC sub-regions in each cluster. All reported p values are exact and two tailed.

## Results

### GFAP+ astrocytes and Iba1+ microglia form the glia limitans externa and neurovascular unit in IC

We first sought to identify the distribution of *GFAP+* astrocytes and *Iba1+* microglia in adult guinea pig IC. Coronal, 60µm sections showed pronounced *GFAP+* and *Iba1*+ labelling of the *glia limitans externa* lining the dorsal and lateral borders of the IC (Figure 1A). Extensive labelling was also distributed medially, lining the cerebral aqueduct, with ramified *GFAP+* astrocytic processes radiating into the periaqueductal grey, as well as the commissure of the IC. Interestingly, we found no *GFAP+* astrocytes throughout the IC parenchyma, save for sparse labelling of cells in the outermost layers of the DCIC and LCIC. Conversely, ramified *Iba1+* microglia tiled the parenchyma in non-overlapping domains with similar density throughout the IC, as quantified in Figure 1B.

**Figure 1.**
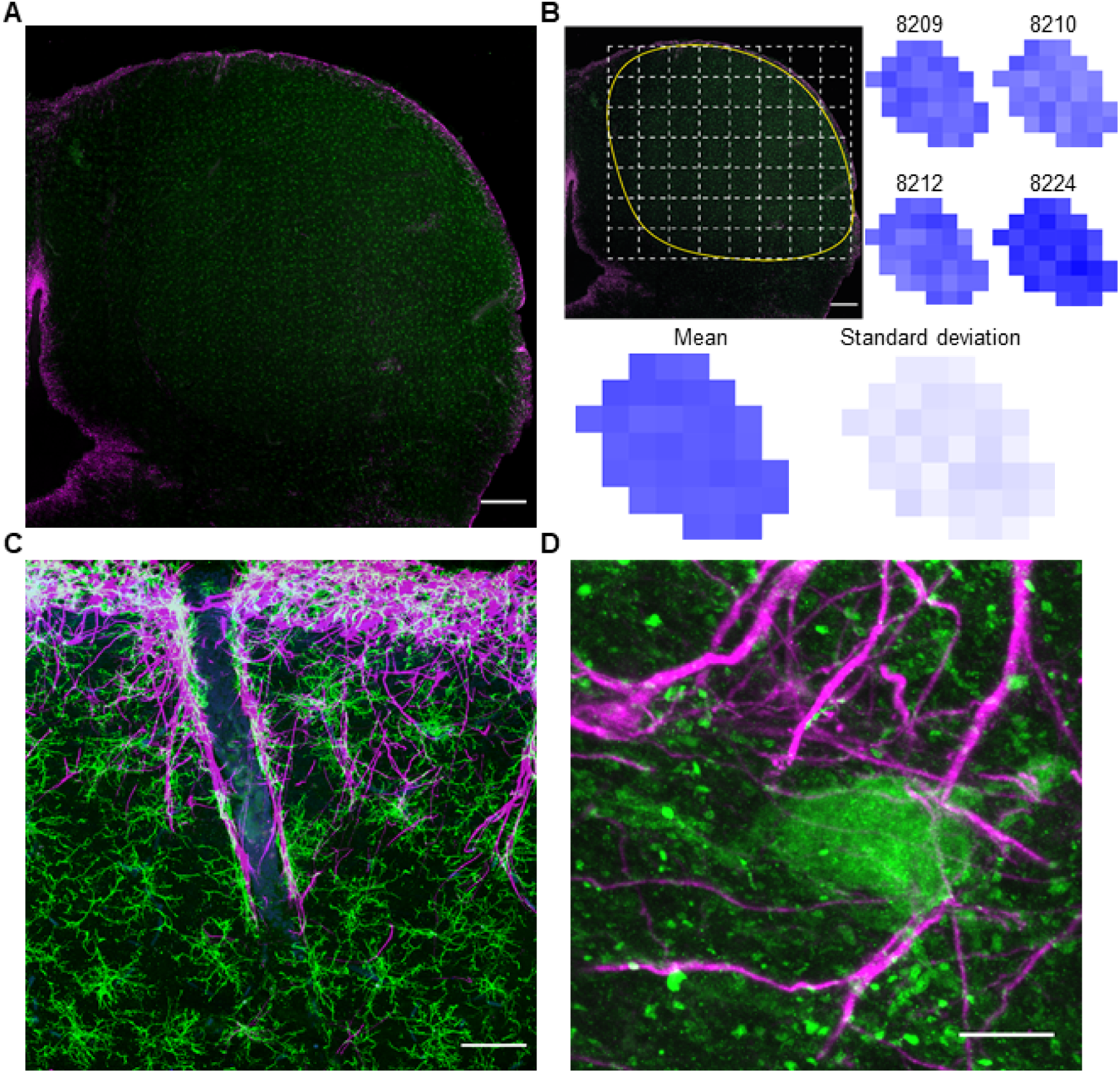
Microglia and astrocytes form the *glia limitans externa* and *peri*-vascular borders but only microglia tile the parenchyma. (A) Tiled maximum intensity projection confocal micrograph of *GFAP+* astrocytes (magenta) and *Iba1+* microglia (green) in IC. Scale bar 400µm. Note that *GFAP* labelling is restricted to the peripheral borders and penetrating vessels while *Iba1* is evenly distributed throughout. (B) Quantification of *Iba1+* somata counts per 450µm^2^ grid (shown inset) within the parenchyma across all four cases. Numbers on top refer to individual animal cases. (C) Confocal micrograph showing *GFAP, Iba1* and *GSL1* (blue) labelling of a penetrating arteriole coursing into IC. Scale bar 50µm. (D) Confocal micrograph showing a *calbindin* (green) expressing neuron in the outer layers of the LCIC surrounded by *GFAP+* axo-somatic processes. Scale bar 10µm.

Combining *Iba1+* and *GFAP+* labelling with the fluorescent-conjugated lectin *GSL1* revealed extensive *peri*-vascular labelling along putative penetrating arteries and arterioles (Figure 1C). Neurons expressing cytoplasmic calbindin or calretinin were distributed in the outermost regions of the cortices of the IC, matching previous reports (Zettel et al., 1997; Ouda et al., 2012) and in close proximity to vessels and *GFAP+* processes (Figure 1D).

These findings demonstrate that many aspects of IC glial organization mirror those reported in other brain regions, with both *GFAP+* astrocytes and *Iba1+* microglia forming the *glia limitans externa* and lining adjacent to blood vessels. However, the observation that *Iba1+* microglia but not *GFAP+* astrocytes were found throughout the parenchyma, suggests a role for *Iba1+* microglia in glial-neuronal putative interactions in IC.

### Distributions of Iba1+ microglia and GAD67+ somata vary between sub-regions of IC

To explore the role of parenchymal microglia in IC, we acquired tiled confocal micrographs of *Iba1+* microglia (Figure 2A) and *GAD67+* neurons (Figure 2B) across whole coronal IC sections (Figure 2C). Labelling revealed putative GABAergic neurons throughout the IC, with increased cell density in high-frequency ventral regions, matching previous reports (Ito et al., 2009; Gleich et al., 2014; Beebe et al., 2016). Dividing the IC into sub-regions based on the criteria of Coote and Rees (2008) allowed definition of ROIs for comparisons between DCIC, LCIC, mid-CNIC and ventral-CNIC (VCNIC), respectively (Figure 2D). Cell counts confirmed a greater number of *GAD67+* neurons in VCNIC (Figure 2E) (H(3)=24.42; p<0.001) than other sub-regions (*post-hoc* Dunn’s tests: VCNIC vs DCIC p<0.0001; VCNIC vs LCIC p=0.0023). Mid-CNIC also had a greater number of *GAD67+* neurons than DCIC (p=0.026) and LCIC (p=0.395). Representative examples of these sub-regional differences are shown in Figure 3.

**Figure 2.**
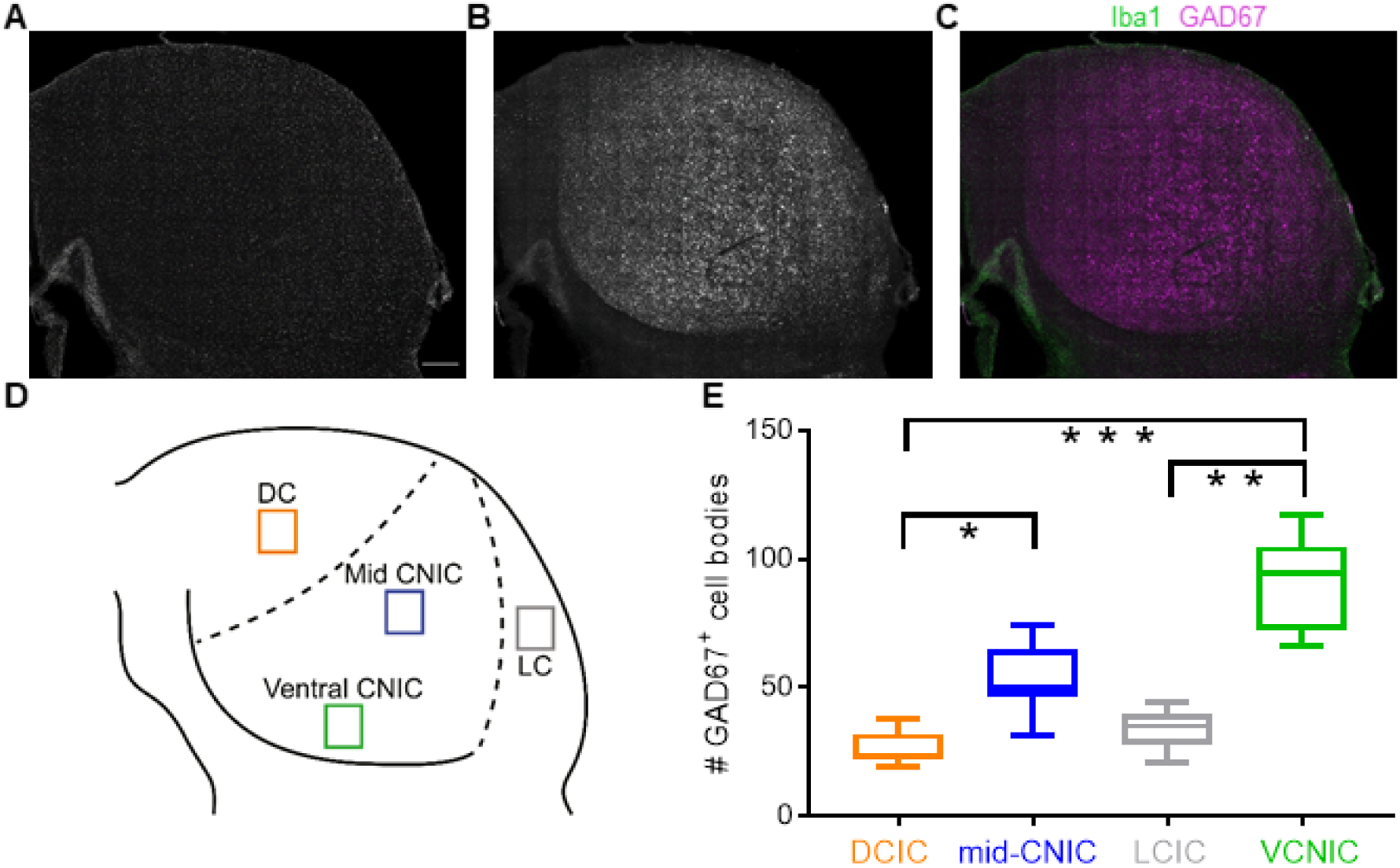
*GAD67+* neurons vary in density between sub-regions of IC. (A) Tiled confocal micrograph showing *Iba1+* microglia tiling IC parenchyma. Scale bar 400µm. Same scale for panels B and C. (B) *GAD67+* neuropil can be seen to demark the medial and ventral borders of the IC. *GAD67+* neurons are found throughout IC but vary in density. (C) Merge of A (green) and B (magenta). (D) Borders of IC sub-regions were delineated using those defined by Coote and Rees (2008). ROIs were located within distinct sub-regions of IC that could be clearly distinguished from one another. (E) Box plot showing *GAD67+* somata counts in each sub-region of interest, across cases. The VCNIC consistently had the highest number of *GAD67+* somata while numbers in DCIC and LCIC were lower than mid-CNIC.

**Figure 3.**
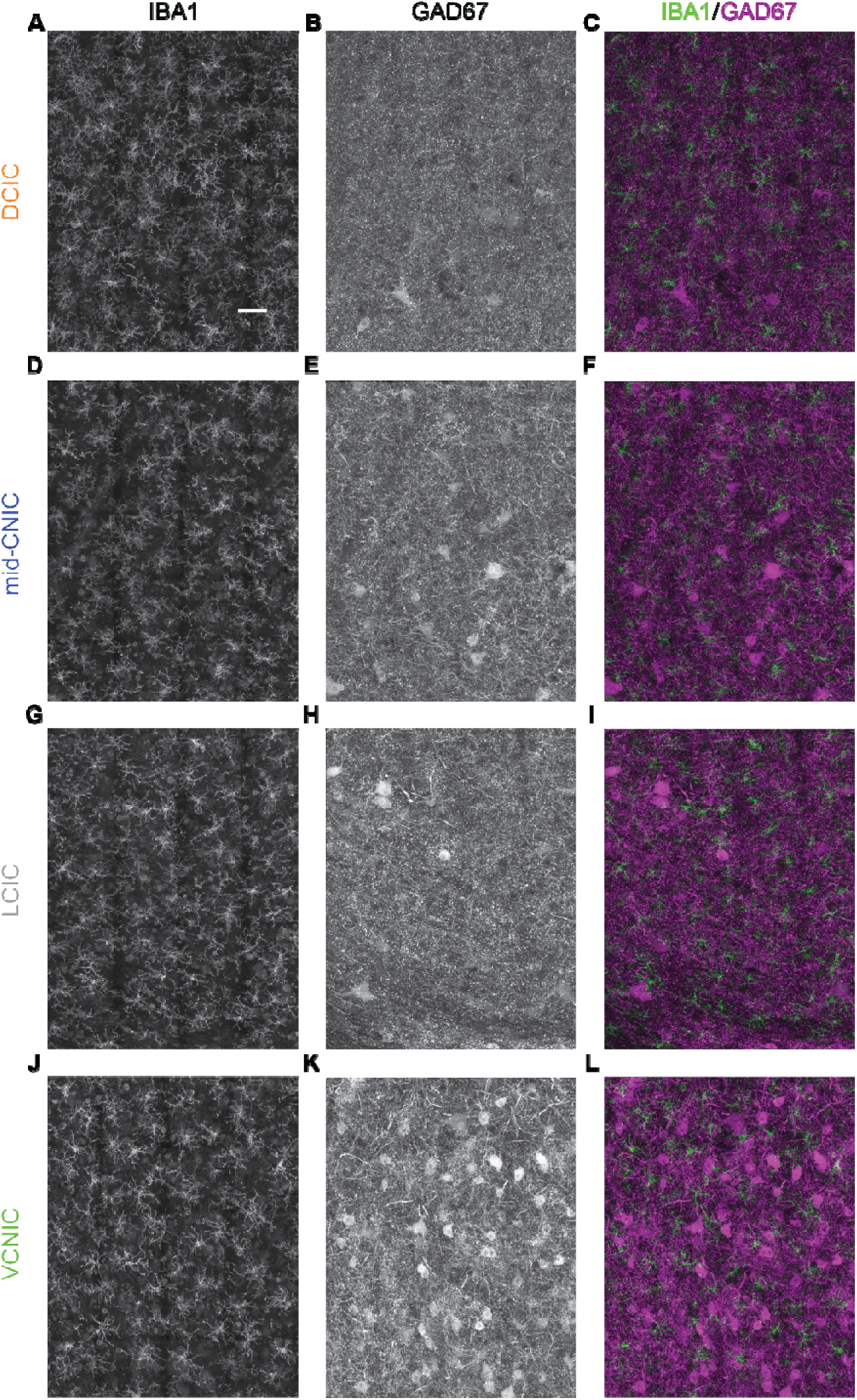
Representative ROI panoramas show differences between *Iba1+* and *GAD67+* cells in sub-regions of IC. Scale bar in (A) 50µm. Same scale for all panels. Maximum intensity projections of tiled confocal panoramas in all sub-regions show (left column; A,D,G,J) *Iba1+* microglia tiling the parenchyma with similar density. Conversely, labelling of *GAD67+* somata (middle column, B,E,H,K) reveals varied cell densities between sub-regions. Merging both labels (right column; C,F,I,L) reveals intercalating of *Iba1+* processes with *GAD67+* labelling.

Analyses of ROIs confirmed similar densities of *Iba1+* microglia cell counts between sub-regions of IC, in spite of the varying density of *GAD67+* cells (Figure 4A). The DCIC had slightly more densely packed *Iba1+* microglia than other sub-regions, with a median of 88 cells (range=70 to 106) per 432×552µm ROI. Mid-CNIC had a median of 83 (range=76 to 95), while LCIC had a median of 80 (range=63 to 104) and VCNIC had a median of 73 (range=57 to 92). However, a Kruskall-Wallis ANOVA with sub-region as the factor found no detectible difference between groups (H(3)=4.91, p=0.179).

**Figure 4.**
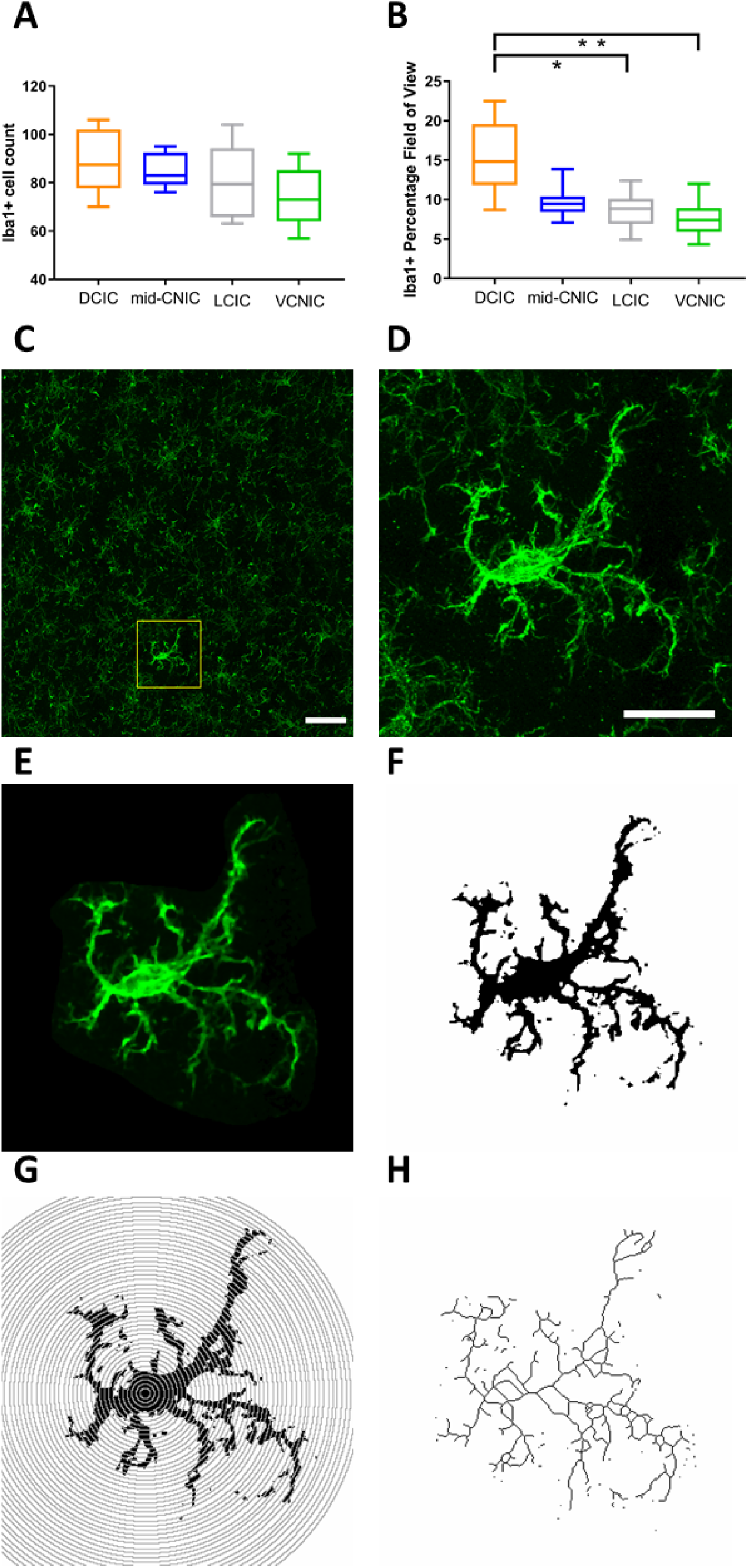
Rationale and process for Sholl and skeleton analyses to quantify morphological characteristics of *Iba1+* microglia. (A) There was considerable overlap in the distributions of *Iba1+* cell counts per ROI between sub-regions, and no statistically detectable differences. (B) Contrastingly, the percentage field of view of *Iba1+* labelling was much greater in DCIC than other sub-regions. These findings suggested a greater number of *Iba1+* process ramifications in DCIC. To test this we therefore undertook morphology analyses as follows: (C) Example of maximum intensity projection of ROI tiled confocal micrographs showing *Iba1+* microglia in IC. A 5-row × 6-column field of view tiled image was taken using 1µm z-slices. Individual cells were subject to Sholl and skeleton analysis (yellow box inset). Scale bar 50µm. (D) Extracted cell from (C) showing high resolution imaging of soma and processes surrounded by non-cellular labelling. Scale bar 20µm. Same scale bar for panels E-H. (E) Non-cellular labelling was cropped and (F) thresholded to binarize cellular processes. (G) Sholl analyses were performed at 1µm resolution from binarized images. (H) Skeletonized cell framework from (F) to be analyzed for number of branches and maximum branch length.

Contrastingly, the percentage field of view of *Iba1+* labelling (above a thresholded binary level, consistent between cases) (Figure 4B) was greater in DCIC (median=14.8%; range=8.7 to 22.5) than mid-CNIC (9.5%; 7.1-13.9), LCIC (8.9% 4.9-12.4) or VCNIC (7.4%; 4.3-12.0). These differences between groups are likely a real effect (H(3)=12.67; p=0.0034; *post-hoc* Dunn’s test between DCIC and VCNIC p=0.002). The greater amount of *Iba1+* labelling in DCIC, despite similar soma density between sub-regions, suggests differences in other aspects of *Iba1+* microglia morphology.

### Iba1+ microglia in DCIC are more ramified than other sub-regions of IC

We predicted that the stronger *Iba1+* microglia labelling in DCIC neuropil was primarily due to a greater number and extent of ramifications compared to other sub-regions of IC. To test this, we conducted Sholl analyses for a total of 64 cells per sub-region (n=256). The maximum intensity projection of each *Iba1+* microglial cell was imaged and analyzed in x and y dimensions.

Cells were identified and selected from ROI images (Figure 4C&D). Background/non-cellular labelling was cropped (Figure 4E) and cellular labelling thresholded to generate binary images (Figure 4F). The number of intersections at every micrometer distance from the center of the soma was calculated (Figure 4G). Binary thresholded cells were also skeletonized to derive information about the shape and structure of ramifications, such as the number of branches and maximum branch length (Figure 4H).

*Iba1+* microglia were more ramified in DCIC than mid-CNIC, LCIC or VCNIC at distances away from the soma (Figure 5A). The total number of intersections of every ramification, across cells (independent of distance from the soma) had a median value of 438 in DCIC (IQR=385 to 500). This was greater than in mid-CNIC (332; ±284-399), LCIC (363; ±326-422), or VCNIC (353; ±302-400) (Figure 5B). A Kruskal-Wallis one-way ANOVA on ranks, with sub-region as the factor found the differences between sub-regions of IC were likely a real effect (H(3)=54.36; p<0.0001). *Post-hoc* analyses via Dunn’s tests showed differences between DCIC and the other three sub-regions were likely to be a real effect (all p<0.0001).

**Figure 5.**
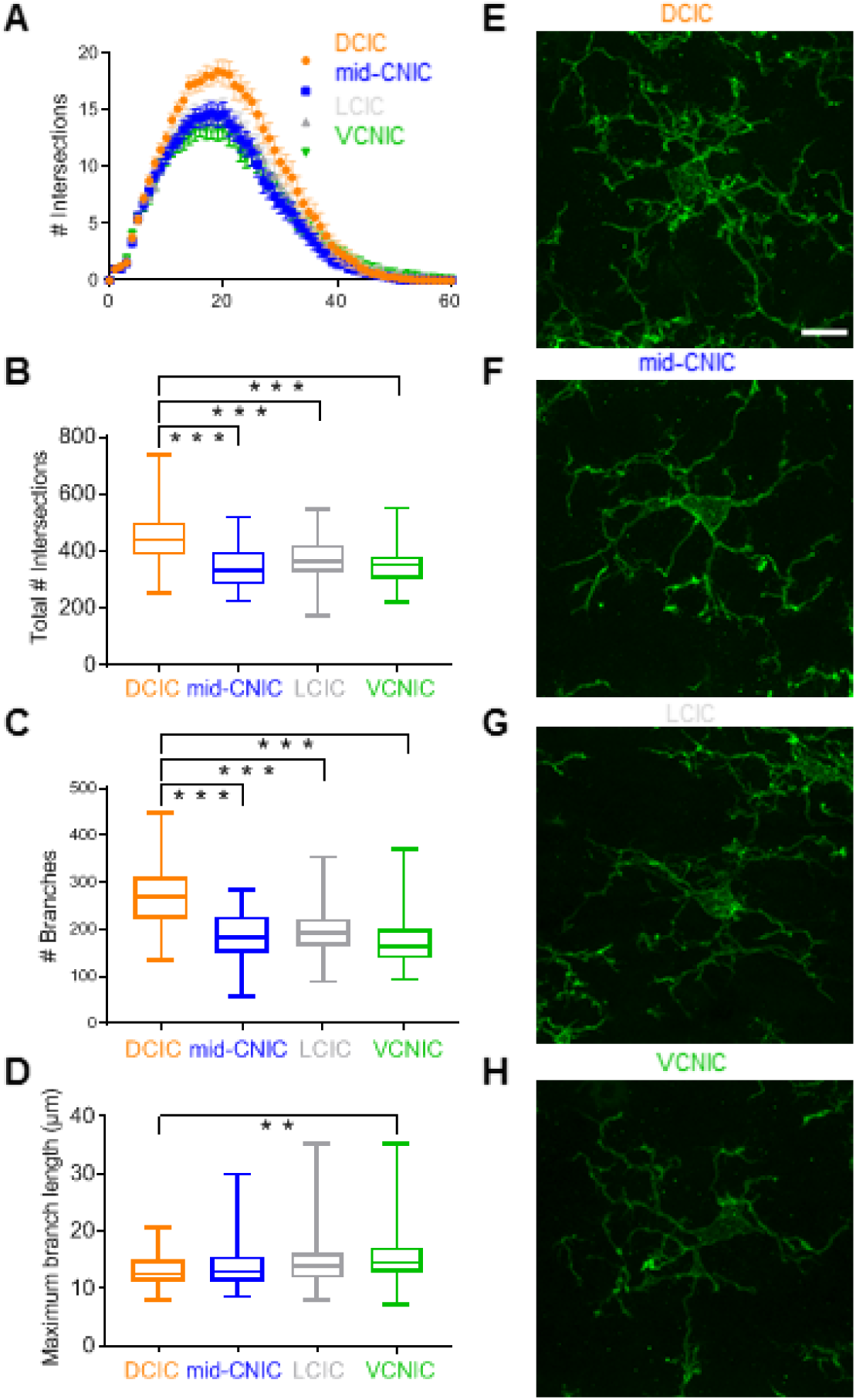
*Iba1+* microglia are more ramified in DCIC than other sub-regions of IC. (A) Sholl analyses (mean ±95% confidence intervals) showing *Iba1+* microglia in DCIC have greater numbers of ramifications than other IC sub-regions at all distances from the soma. (B) Total number of ramification intersections independent of distance from soma are greater in DCIC. (C) Greater number of intersections in DCIC are due to a greater number of branching ramifications. (D) Maximum branch length, a measure of how long ramifications travel before branching, are longest in VCNIC and shortest in DCIC. Representative examples of *Iba1+* microglia in (E) DCIC, (F) mid-CNIC, (G) LCIC and (H) VCNIC. Scale bar in (E) 10µm. Same scale bar for panels (F-H).

Analyses of skeletonized *Iba1+* microglia revealed those in DCIC also had a greater number of branches (Figure 5C), with a median of 269 (±222-312). This was greater than mid-CNIC (183; ±150-229), LCIC (193; ±165-224), and VCNIC (165; ±138-201). These differences were also likely a real effect (H(3)=71.30, p<0.0001; Dunn’s tests DCIC vs other sub-regions all p<0.0001).

Conversely, maximum branch length, defined as the longest distance covered by any ramification of skeletonized *Iba1+* microglia without branching, followed the opposite trend. Longest maximum branch lengths were found in VCNIC (median=14.44µm; IQR=12.76-17.18) (Figure 5D). Shorter distances were found in LCIC (14.01; IQR±=11.92-16.16), mid-CNIC (12.86; ±11.22-15.60) and DCIC (11.62; ±11.26-15.01). The difference between VCNIC and DCIC was likely a real effect (H(3) =12.18, P = 0.0068; *post-hoc* Dunn’s test p=0.0079). Representative examples of *Iba1+* microglia in each sub-region of IC are shown in Figure panels 5E-H.

### Iba1+ putative abutments reveal two novel types of GAD67+ neurons in IC

We hypothesized that *GAD67+* neurons, which are known to receive a variety of types of presynaptic contacts (Ito et al., 2009; Beebe et al., 2016), may also receive different types of *Iba1+* abutments. To examine the nature of *Iba1+* putative interactions with *GAD67+* neurons, we quantified five dependent variables from each cell ROI (n=40 per sub-region): (i) *GAD67+* soma maximum diameter; (ii) percentage of *GAD67+* soma abutted by *Iba1+* processes; (iii) number of *Iba1+* microglia with processes abutting each *GAD67+* soma; (iv) number of distinct *Iba1+* processes abutting each *GAD67+* soma; and (v) total length of *Iba1+* processes abutting each *GAD67+* soma (µm). The methods used to calculate these variables are shown in Figure 6. These features were calculated from micrographs such as the representative example in Figure 7A, which shows a *GAD67+* neuron being abutted by two *Iba1+* microglia. A correlation matrix revealed weak associations between *GAD67+* soma maximum diameter (i) and the other four dependent variables (ii-v) (Figure 7B). There were stronger correlations between the four *Iba1* related variables (ii-v). As these variables were only weakly correlated with *GAD67+* neuron diameter, we further investigated whether a multivariate analysis could better explain the observed distributions.

**Figure 6.**
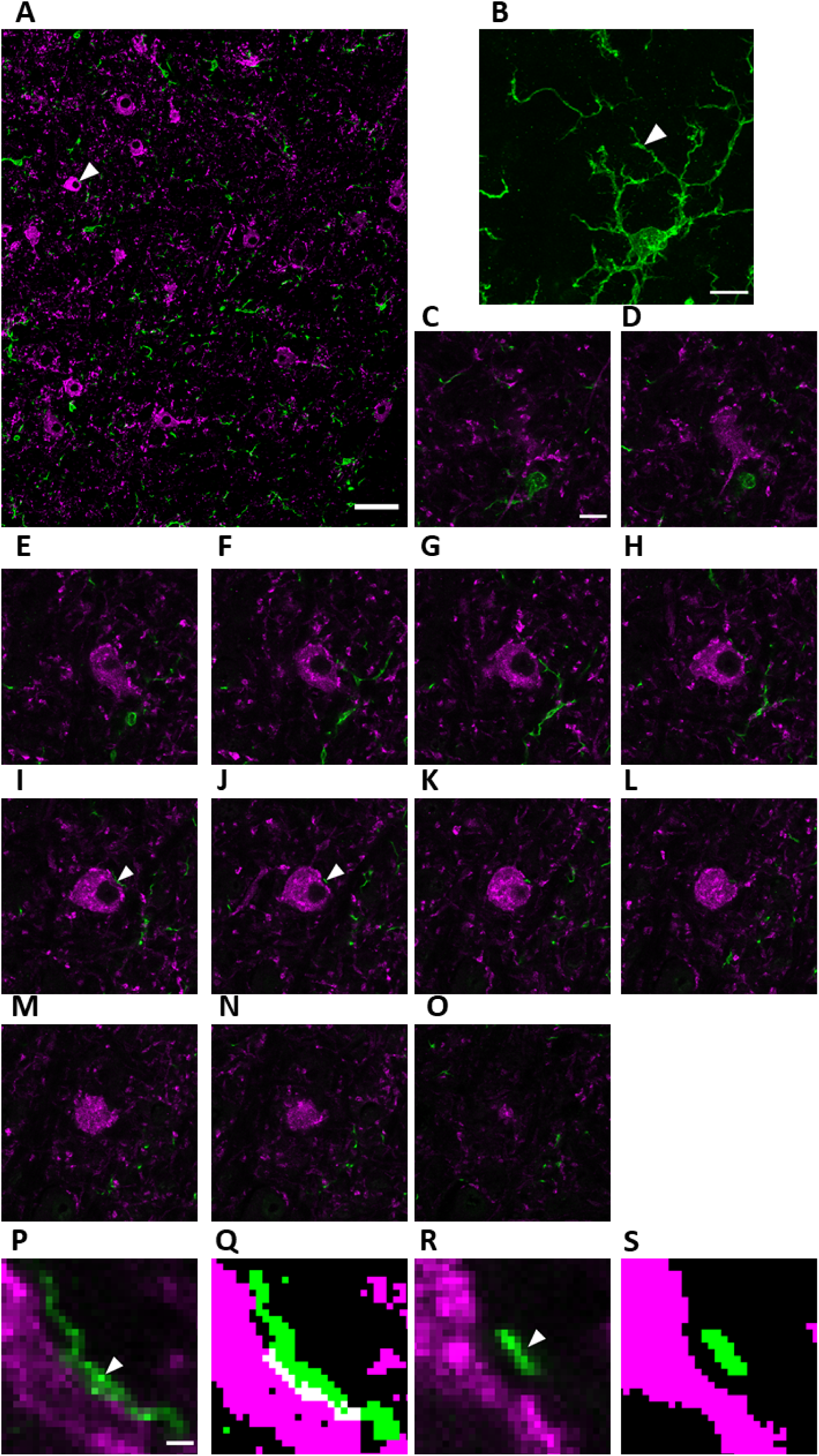
*GAD67+* cells were selected that were contained entirely within the x, y and z coordinates of the tissue section and ROI. (A) Example 1µm slice of an ROI panorama in VCNIC, labelled for *GAD67* (magenta) and *Iba1* (green). Example *GAD67+* cell (white arrow) was selected for analysis. Scale bar 50µm. (B) Maximum intensity projection of *Iba1+* cell through 13 z-planes in zoomed region around cell selected in (A), allowing visualization of microglial processes and soma. Scale bar 10µm. (C-O) Example 1µm slices though rostro-caudal axis in the z-plane. Soma perimeter length was measured in each slice. White arrows in (I&J) point to magnified region in (P-S). Scale bar 10µm. (P) Magnified region from (I) demonstrating an *Iba1+* process abutting *GAD67+* soma. Scale bar 1µm also applies to panels (Q-S). (Q) Thresholding using the Huang algorithm in ImageJ for each label reveals abutment of *Iba1+* process onto *GAD67+* soma, with no pixels separation, and a 1-2 pixel thick plexus of double labelling (white), allowing quantification of the location and length of abutment. The distance of abutment was measured along the *GAD67+* perimeter. Each distinct process abutting the *GAD67+* cell was counted and the total number of *Iba1+* cells abutting the *GAD67+* cell were measured. (R) Magnified region from (J), showing *Iba1+* process in close proximity to but not abutting GAD67+ soma. Thresholding each label reveals clear separation with unlabeled pixels between. This demonstrates that the measures taken facilitated positive and negative identification of abutments at 1µm intervals through the z-plane of all cells analysed.

**Figure 7.**
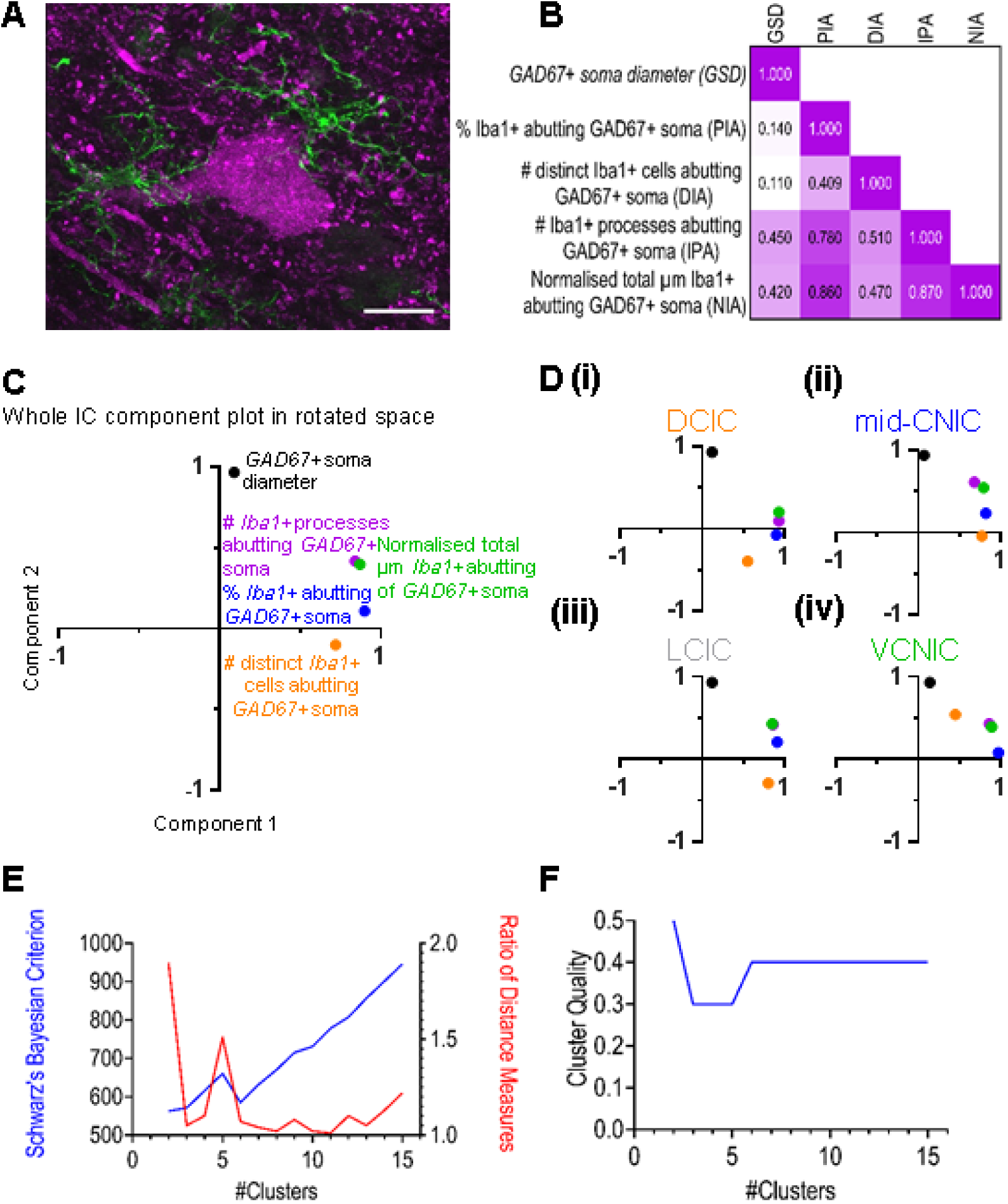
(A) Representative *GAD67+* cell (magenta) receiving abutting somatic processes from two *Iba1+* microglia (green). Scale bar 20µm. (B) Correlation matrix showing Spearman’s rank correlation coefficient for each combination of measures derived from putative interactions between *Iba1+* microglia and *GAD67+* cells. Note lower values in first column. (C) Principal component analysis of the five variables in (B). *GAD67+* soma diameter separated from the other four variables, which grouped together. (D) As (C) but for each sub-region in IC, showing similar findings in all sub-regions, suggesting robust clustering throughout IC. (E) The largest difference between Schwarz’s Bayesian criterion and the ratio of Euclidean distance measures is observed in cluster 2, demonstrating that two clusters displays the best explanatory power. (F) Silhouette measures of cohesion and separation were compared from 2-15 clusters, with two clusters demonstrating the highest cluster quality. These data demonstrate that not only did two clusters exhibit the best available explanatory power, but also strong clustering quality, reflecting real underlying differences in *Iba1+* abutments onto *GAD67+* neurons between clusters.

We conducted a principal component analysis for the five variables in all sub-regions of the IC. The data showed a clear dissociation between *GAD67+* neuron diameter in one cluster and the other four variables, which clustered together (Figure 7C). Both clusters were categorized using a standard correlation coefficient of >0.5 as a cut-off value, which showed one cluster was explained by only the *GAD67+* neuron diameter variable, while the other cluster had significant contributions from all four of the *Iba1+* related variables. These trends were also true for all sub-region specific analyses in IC (Figure 7Di-iv). We then conducted a two-step cluster analysis including all five variables. We employed Euclidean distance measures with Schwarz’s Bayesian clustering criterion (Figure 7E). Contrasting iterations up to 15 clusters, the analysis found two clusters demonstrated the best explanatory power, displaying good (0.5) silhouette measures of cohesion and separation (Figure 7F).

All 160 cell ROIs were classified into one of the two clusters determined by the two-step analysis. Representative examples of each cluster found in each sub-region are shown in Figure 8. There were 23 cases in cluster 1, and 137 in cluster 2. To visualize the contribution of each of the four *Iba1* related variables, each was plotted as a function of *GAD67+* neuron diameter (Figure 9A-D). These scatterplots revealed a dissociation with little overlap between the two clusters using the percentage *Iba1+* abutting *GAD67+* somata (Figure 9A). There was almost perfect discrimination between the clusters of the normalized total µms of *Iba1+* abutments onto *GAD67+* neuron somata (Figure 9B). Conversely, the number of *Iba1+* cells abutting each *GAD67+* soma had a significant degree of overlap with little difference between clusters (Figure 9C). The number of *Iba1+* processes abutting *GAD67+* somata had little overlap between distributions (Figure 9D).

**Figure 8.**
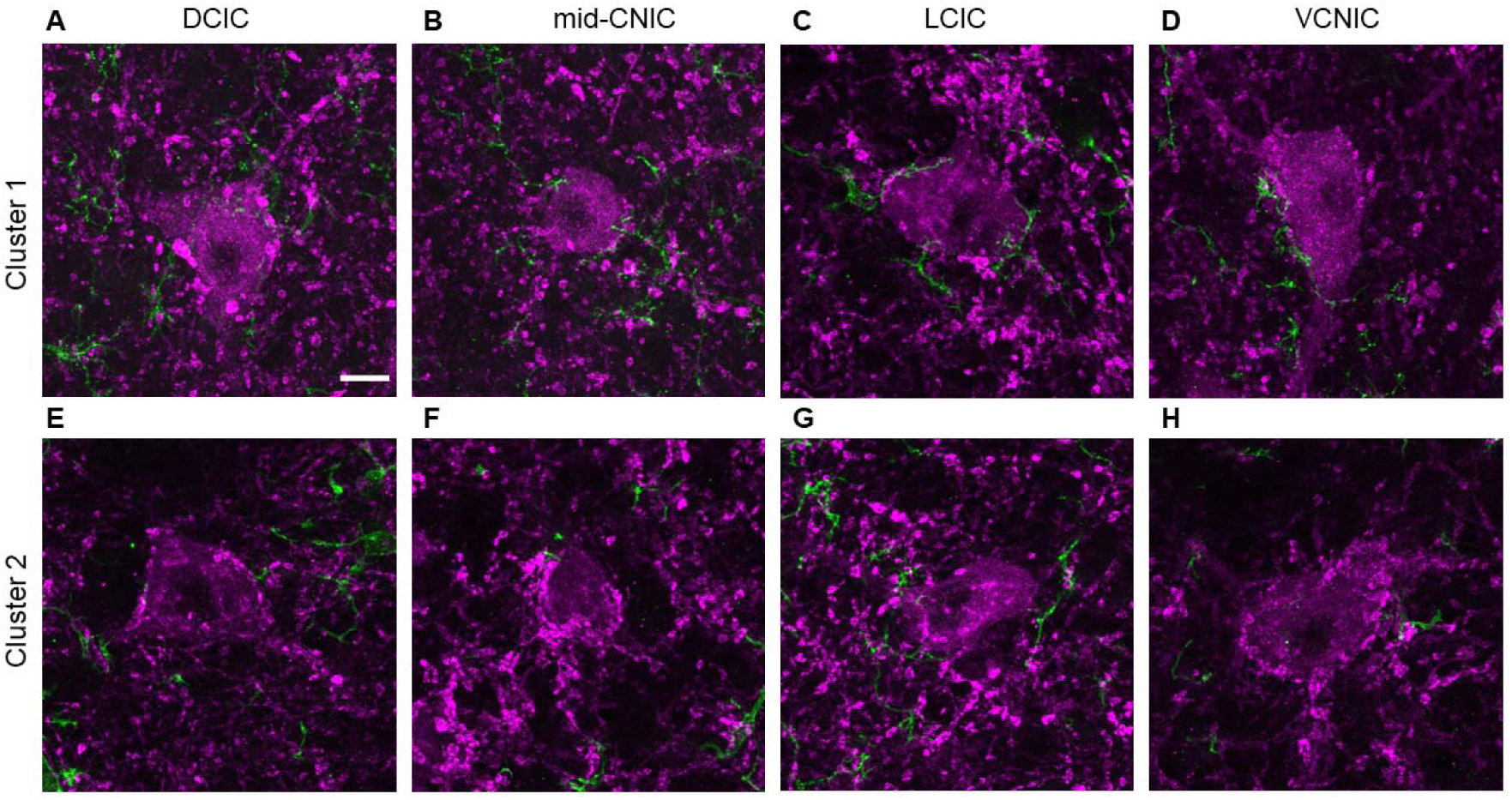
Example *GAD67+* cells taken from each sub-region of IC (maximum intensity projection of five 1µm slices through the center of each cell in the rostro-caudal z-plane), with one example from each designated cluster from the two-step analysis. Scale bar 10µm applies to all images. Note the greater length of *Iba1+* abutments surrounding *GAD67+* somata in cluster 1 (top row), while adjacent Iba1+ processes in cluster 2 make few abutments onto the somata (bottom row).

**Figure 9.**
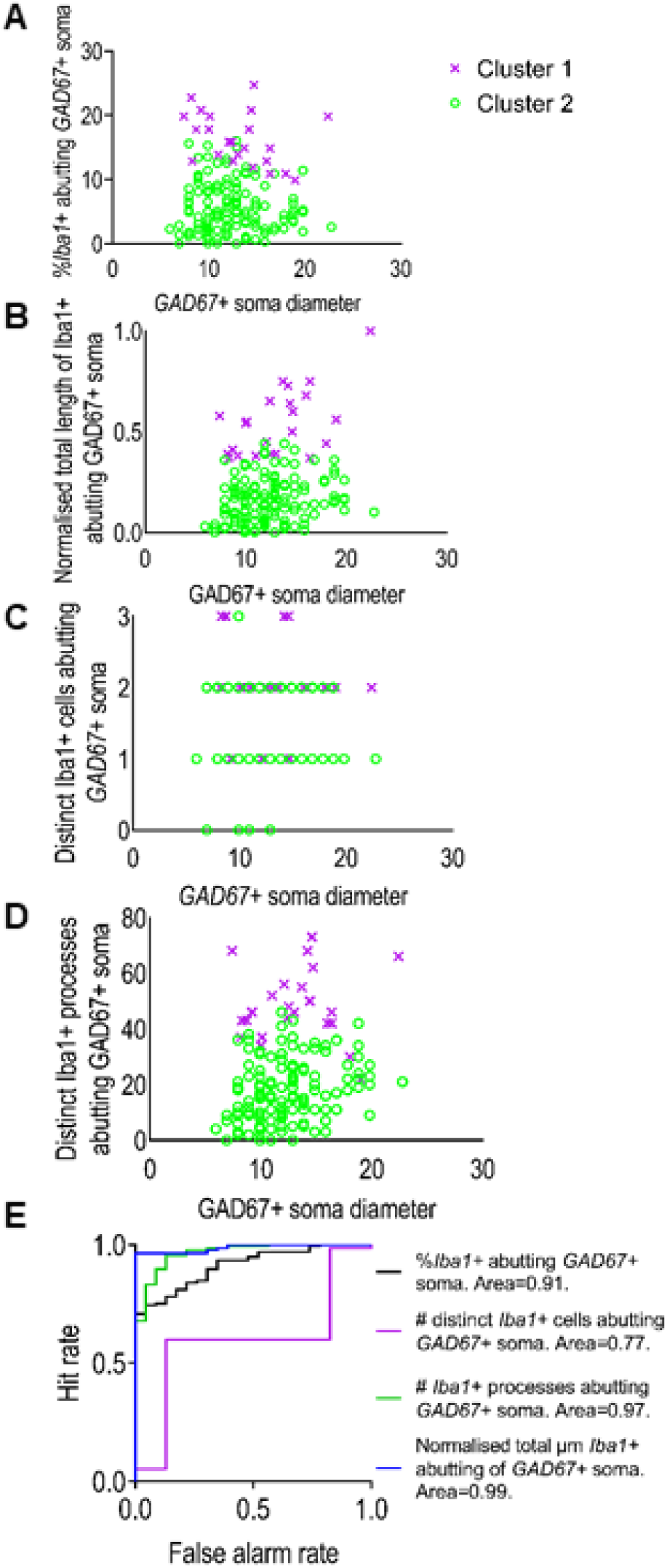
*GAD67+* neurons in IC can be classified into two clusters based on the amount of somatic *Iba1+* abutments they receive. Scatterplots showing each *GAD67+* cell from the cluster analysis, classified into either cluster 1 (magenta crosses; n=23) or cluster 2 (green open circles; n=137) for each *Iba1+* related variable, plotted as a function of *GAD67+* soma diameter: (A) Percentage *Iba1+* labelling abutting *GAD67+* somata; (B) normalized total length of *Iba1+* labelling abutting *GAD67+* somata; (C) distinct number of *Iba1+* cells abutting *GAD67+* somata; (D) number of *Iba1+* processes abutting *GAD67+* somata. (E) ROC analyses showing classifier performance of each variable in discriminating *GAD67+* cells into cluster 1 or cluster 2. Normalized total µm *Iba1+* abutting and number of processes of *GAD67+* somata could almost perfectly discriminate between clusters.

To compare the ability of each of these variables to independently discriminate between the two clusters, we conducted ROC analyses (Figure 9E). These data revealed that while each variable had an area under the curve >0.5, the three variables relating to the nature of *Iba1+* processes abutting *GAD67+* neurons had the best discriminatory power.

### Iba1+ putative interactions with GAD67+ neurons show little difference between sub-regions of IC

We explored whether any of the variables or clusters identified had a relationship to the sub-region of IC in which the cells were located. Diameter of *GAD67+* somata did not vary between sub-regions (Figure 10A) (H(3)=5.3; p=0.151). A small difference was found for the percentage of *GAD67+* soma abutted by *Iba1+* processes (Figure 10B) (H(3)=9.9; p=0.019). *Post-hoc* Dunn’s tests suggested a potential real difference between DCIC and VCNIC (p=0.012). However, there was extensive overlap between the distributions so there is a reasonable chance this may not be a true effect. The number of *Iba1+* cells abutting each *GAD67+* somata (Figure 10C) (H(3)=6.44; p=0.092), the number of *Iba1+* abutments onto *GAD67+* somata (Figure 10D) (H(3)=5.55; p=0.141), and the normalized total number of µm covered by *Iba1+* abutments onto *GAD67+* somata (Figure 10E) (H(3)=4.64; p=0.200) did not differ between sub-regions. However, the number of *GAD67+* cells abutted by each *Iba1+* cell was greater in VCNIC (Figure 10F) (H(3)=21.32; p=0.0006). *Post-hoc* Dunn’s tests revealed a likely real difference between VCNIC and DCIC (p=0.005). We interpret this as being due to the greater density of *GAD67+* neurons in VCNIC (Figure 2E).

**Figure 10.**
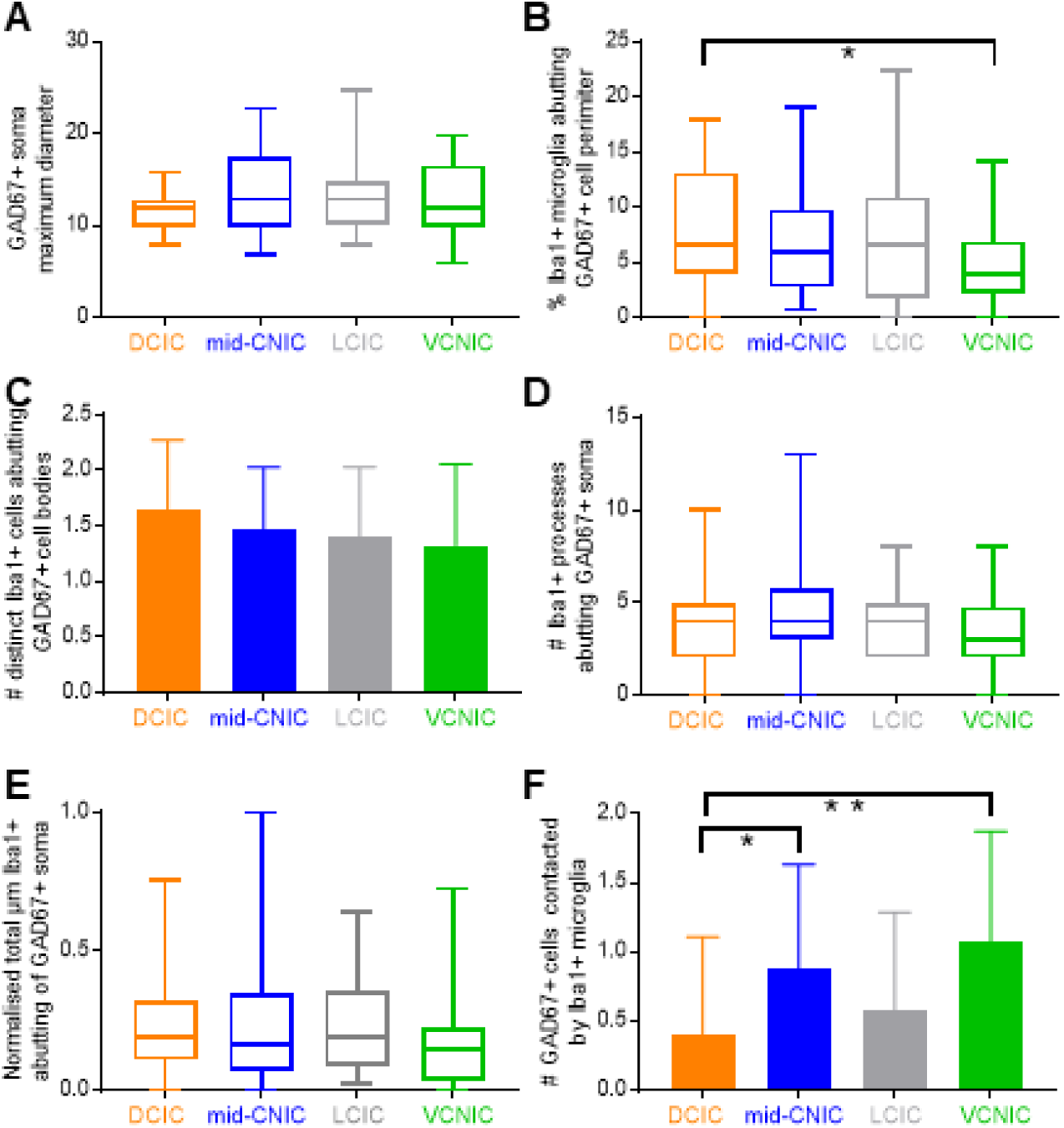
No clear relationship between measured variables and IC sub-region. (A) *GAD67+* soma diameter was similar across sub-regions of IC. (B) The percentage of *Iba1+* abutting *GAD67+* somata was lower in VCNIC than the other three sub-regions. (C) The number of *Iba1+* cells abutting *GAD67+* somata, (D) the number of *Iba1+* processes abutting *GAD67+* somata, and (E) the normalized total µm of *Iba1+* abutting of *GAD67+* somata did not differ between sub-regions. (F) There were, on average, a greater number of *GAD67+* somata abutted by each *Iba1+* cell, owing to the greater density of *GAD67+* cells in VCNIC.

Both clusters had similar proportions of ROIs from each of the four sub-regions of IC (cluster 1: 7 DCIC (30%), 8 mid-CNIC (35%), 6 LCIC (26%), 2 VCNIC (9%); cluster 2: 33 DCIC (24%), 32 mid-CNIC (23%), 34 LCIC (25%), 38 VCNIC (28%)) A Chi-squared test suggested that there was no difference in the relative proportion of each cluster, between sub-regions (χ^2^(3)=4.21 p=0.24).

## Discussion

These findings reveal that *Iba1+* microglia, but not those of *GFAP+* astrocytes, are present throughout the parenchyma of the IC in the healthy, mature auditory system. Taking advantage of the specialized and functionally diverse sub-regions of IC, we found the first evidence, to our knowledge, revealing differences in microglial morphology, between sub-regions. Interestingly, *Iba1+* microglia in DCIC, a sub-region known to receive a greater proportion of glutamatergic corticofugal contacts, are more ramified than those in other sub-regions of IC. We also developed a new analysis method, which was used to investigate the number and length of abutting microglial processes onto *GAD67+* neuronal somata. Multivariate cluster analyses were applied to semi-quantify variables of abutments and revealed two distinct types of *GAD67+* neuron in IC, distinguished by the extent of *Iba1+* microglial abutments onto their somata. Taken together, these data demonstrate that microglia in IC exhibit anatomical features and connectivity with GABAergic neurons suggesting specialization of function relating to the local demands of processing.

### Significance of IC sub-regional differences in microglial morphology

Sensory processing can be interpreted through triad models of organization, based on observations that central sub-regions of sensory pathways are dominated by ascending innervation, producing brisk responses at short latencies to simple stimuli, such as in the CNIC. Across different sensory nuclei, there are at least two sub-regions located adjacent to central sub-regions, which typically require more complex stimuli to elicit responses and which occur at longer response latencies. One of these sub-regions typically receives a diversity of polymodal inputs (LCIC), while the other is primarily driven by descending, top down afferents (DCIC). These sub-regions exhibit differing connectional as well as cytoarchitectonic and chemoarchitectonic organization (Sweet et al., 2005). In IC, there is longstanding evidence for a triad model of organization using a diversity of methodologies, including electrophysiology (Syka et al., 2000), fMRI (Baumann et al., 2011; De Martino et al., 2013), histology (Faye-Lund and Osen, 1985), immunohistochemistry (Coote and Rees, 2008) and tract tracing of projections (see Malmierca and Hackett (2010) and Schofield (2010)). The present study provides evidence that *Iba1+* microglia morphologies also exhibit differences between sub-regions (Figure 5).

Why might microglia exhibit differing morphologies in distinct sub-regions of IC? There is longstanding evidence that microglia are sensitive to their local environment and exhibit morphological differences throughout the brain (Lawson et al., 1990). There are clear neuroanatomical differences between CNIC, DCIC and LCIC, including increased cytoarchitectural and myeloarchitectural density in CNIC (Faye-Lund and Osen, 1985). There is also a more defined laminar organization of the CNIC than the DCIC, in part due to the neuronal morphologies present as well as coursing fibres (Oliver and Morest, 1984). As the DCIC is known to have larger neuron sizes than CNIC, one might speculate that the increased number of branching ramifications we observed in DCIC (Figure 5) correlates with larger neuronal somata. However, the LCIC is also known to have larger neurons than CNIC, but had a similar number of microglial branching ramifications to mid-CNIC and VCNIC. This suggests that differing microglial morphologies may relate to other aspects of local processing. For instance, it is known that microglia interact at synapses (Trapp et al., 2007; Wake et al., 2009; Tremblay et al., 2010). The greater number of branching ramifications in DCIC may therefore relate to the differing nature of local synaptic processing therein compared to CNIC and LCIC. Certainly, cortical regions of IC exhibit much stronger novelty detection and stimulus specific adaptation than CNIC (Ayala and Malmierca, 2013). This may partly relate to the primary afferent drive to DCIC being descending corticofugal fibers (Herbert et al., 1991; Winer et al., 1998; Bajo and Moore, 2005; Bajo et al., 2006). Projections to DCIC from auditory cortex originate from glutamatergic (Feliciano and Potashner, 1995) pyramidal cells in layer V (Games and Winer, 1988; Winer and Prieto, 2001). Corticofugal inputs to DCIC primarily target glutamatergic IC neurons (Nakamoto et al., 2013), while large *GAD67+* neurons in DCIC are a source of tectothalamic inhibition (Geis and Borst, 2013). The DCIC receives a greater diversity of inputs than CNIC, including from auditory cortex, the contralateral IC (Orton et al., 2016), and ascending projections from brainstem. Corticofugal inputs to DCIC have been shown to play an essential role in auditory learning and plastic reweighting of cues (Bajo et al., 2010; Keating et al., 2013). These connections likely underlie elements of human adaptation in unilateral hearing loss (Kumpik and King, 2019), though molecular mechanisms for these observations are underexplored. Microglial influence over these various aspects of auditory processing remains to be explored.

Another explanation may be the higher density of neurons in CNIC. There has been previous suggestion that microglial cell density and neuronal cell density are inversely related (Lawson et al., 1990). However, we found no strong evidence for differences in microglial cell density between IC sub-regions using two different levels of approach (Figure 1B&4A). This was in spite of much greater numbers of *GAD67+* somata in VCNIC than other sub-regions (Figure 2&3). A third possibility is that there are relationships with the myeloarchitectural differences between sub-regions, however, DCIC and VCNIC exhibit similar, high levels of axons of passage compared to mid-CNIC, arguing against such an explanation. We therefore have no strong evidence as to why microglia in DCIC are more ramified that other sub-regions and this requires further investigation.

### Two novel clusters of GABAergic neurons

As the auditory pathway contains a large proportion of inhibitory neurons, with around a quarter of neurons being GABAergic (Oliver et al., 1994; Merchán et al., 2005), understanding their structure, function, and organization is a question of fundamental importance. Previous approaches to classifying *GAD67+* neurons in IC have focused on soma size (Roberts and Ribak, 1987a, 1987b; Ono et al., 2005) coupled with axo-somatic inputs (Ito et al., 2009), perineuronal nets (Beebe et al., 2016) or cytoplasmic calcium binding protein expression (Ouda and Syka, 2012; Engle et al., 2014). The present analyses show that while there is merit to these approaches, other features of GABAergic sub-types exist. Indeed, we have discovered that *GAD67+* cells can be classified into two distinct clusters based on the total amount of *Iba1+* abutments onto their soma (Figures 7,8&9). That GABAergic neurons in IC can be defined based on *Iba1+* inputs suggests that microglia are essential to the structure and function of GABAergic processing in the mature, adult auditory system.

It may be claimed that cluster analyses, as well as other classification approaches, can produce artificial discriminations between data that reside along continua, such as has been reported for frequency response areas in IC (Palmer et al., 2013). Indeed in many cases, forcing data through cluster analyses will produce clustered data, irrespective of whether these clusters represent meaningful differences. However, we permuted our cluster analysis through various iterations and found that not only could the data be best clustered by two clusters, but that those clusters had strong explanatory power, with good silhouette measures of cohesion and separation and good cluster quality (Figure 7E&F). Furthermore, visualization of representative examples of these clusters showed that those cells in cluster 1 clearly received a greater number and length of abutting *Iba1+* processes onto their somata than those in cluster 2 (Figure 8).

ROC analyses revealed that the two clusters could be distinguished by three variables that quantified aspects of *Iba1+* processes, but not by the number of *Iba1+* microglial cells abutting each *GAD67+* neuron. This may reflect the highly motile and dynamic nature of microglial processes (Wake et al., 2009). Other features of GABAergic neurons in IC, such as their discharge patterns and expression of associated ion channels also do not relate to soma size (Ono et al., 2005). Interestingly, the two identified clusters of *GAD67+* neurons did not differ in their relative proportion between the four sub-regions in IC. Future work may investigate the differing afferent neural inputs to and efferent targets of these cells, to identify likely physiological and connectional differences between clusters and their relationship to GABAergic processing.

### Technical Considerations

The use of primary antibodies in less studied species such as guinea pig can be challenging due to potential differences in epitopes and when not adequately controlled for, may lead to spurious observations (Schonbrunn, 2014). This is important when using exploratory approaches as in the present study, to ensure all analyses are predicated on specific and selective labelling (Voskuil, 2017). We therefore conducted extensive control experiments, excluding primary antibody only, secondary antibody only and both antibodies, to ensure analyses were based on true labelling.

The lack of *GFAP+* astrocytes in IC parenchyma (Figure 1) was surprising and necessitated confirmatory experiments. However, in all cases, a lack of *GFAP+* astrocytes in the parenchyma was found alongside extensive labelling in *peri*-vascular regions and the *glia limitans externa*, demonstrating consistency within and between cases. The lack of *GFAP+* astrocytes in IC parenchyma does not exclude the possibility that astrocytes reside throughout IC. Indeed, a recent report employing SR101 revealed a network of putative astrocytes throughout CNIC (Ghirardini et al., 2018). However, there is some labelling of oligodendrocytes with this marker, which hampers interpretability in studies trying to selectively label astrocytes (Hill and Grutzendler, 2014). There are a diversity of other non-GFAP markers that may reveal the distribution of distinct astrocyte sub-types throughout the IC, including the parenchyma, however, this was beyond the scope of the present study.

Functional differences between astrocytes in CNIC and the outer layers of DCIC and LCIC have been suggested previously via 3-chloropropanediol-induced lesions, which selectively destroyed the former but not the latter (Willis et al., 2003; Willis et al., 2004). The present study leads to the speculation of fundamental gliochemical and physiological differences that may relate to the sub-region specific roles microglia and astrocytes play in their local milieux (Lawson et al., 1990; Olah et al., 2011). Recently, RT-PCR of single IC astrocytes revealed expression of functional inhibitory neurotransmitter transporters *GlyT1*, *GAT-1*, and *GAT-3* (Ghirardini et al., 2018). Sub-regional differences in *GAD67+* neurons in the present study suggest that GABAergic and glycinergic signaling released from and received by glial cells may also exhibit such variations throughout IC and perhaps in other structures.

### Conclusions

We have described, for the first time, that *Iba1+* microglia, but not *GFAP+* astrocytes tile the adult IC parenchyma and have discovered sub-regional differences in the morphology of microglia in IC. Furthermore, multivariate statistical approaches revealed two new clusters of *GAD67+* neurons which can be distinguished based on the total amount of *Iba1+* abutments they receive from microglial processes. Our findings demonstrate morphological (and suggest putative functional) diversity amongst IC microglia, with differential ability to interact with GAD67+ somata. These data highlight the fundamental role microglia play in the organization and likely function of sensory systems in the healthy, mature brain.

## Acknowledgements

We thank Adrian Rees for generous donation of tissues and Claudia Racca for comments on an earlier version of the manuscript.

## Abbreviations

IC: inferior colliculi
CNIC: central nucleus of the inferior colliculi
DCIC: dorsal cortex of the inferior colliculi
LCIC: lateral cortex of the inferior colliculi
VCNIC: ventral central nucleus of the inferior colliculi
Iba1: ionized calcium binding adaptor molecule 1
GAD67: glutamic acid decarboxylase
PBS: phosphate buffered saline
GFAP: glial fibrillary acidic protein
ROI: region of interest
ROC: receiver operating characteristic

